# Recovery of the type specimen of *Avena breviaristata*, an endemic Algerian grass species collected only once (1882): morphology, taxonomy and botanical history

**DOI:** 10.1101/691717

**Authors:** Jennifer Gabriel, Natalia Tkach, Martin Röser

## Abstract

*Avena breviaristata*, collected only once (1882) in Algeria and never re-collected since, is a very mysterious grass species because unfortunately even the type specimen got lost 60--80 years ago. Morphological information on this species was thus based on a few published descriptions, which made it difficult, however, to correctly infer the genus affiliation of this morphologically odd species. *Avena breviaristata* became affiliated in the past with various oat-like genera (*Avenula*, *Helictotrichon*, *Tricholemma*). Due to the recent rediscovery of the type specimen at the P herbarium and the opportunity to study this specimen, we report here on the morphological characters of *A. breviaristata* underpinned by meaningful illustrations. They are discussed in comparison with the morphology of representative species of the above-mentioned genera. Uncommon characters of the spikelets (type of disarticulation of the rachilla, lemma structure, lodicules) and to some extent of the inflorescences, leaves and leaf sheaths support the inclusion of *A. breviaristata* in the North African genus *Tricholemma*. Considering biogeography, *T. breviaristatum* from the arid Hauts Plateaux in Algeria is a highly xeromorphic counterpart of the mesomorphic species *T. jahandiezii*, which is confined to higher altitudes of the rather humid Moyen Atlas in Morocco. This underlines the status of *Tricholemma* as a relic endemic. Our morphological survey supports the classification of *Avenula* (only *A. pubescens*) as separate from *Helictotrichon* s.str. and *Helictochloa*. Moreover, morphological evidence does not support an origin of *A. pubescens* by intergeneric hybridization between the latter genera as hypothesized in some prior studies. Especially the glabrous palea, the special shape of the lodicules and the structure of the awn show no intermediacy. The complicated history of the type collection of *T. breviaristatum* and the role of botanical authors are given.

## INTRODUCTION

The Mediterranean is one of the hotspots of global plant biodiversity (Myers et al., 2000). Apart from comparatively widespread species in this region (circum-Mediterranean) there are many narrowly distributed taxa, which are characteristic of partial areas, frequently confined to the eastern or western, more rarely to the southern Mediterranean (Médail & Quézél, 1997; Thompson, 2005; Médail & Diadema, 2009). This is at least partly due to the comparatively narrow overall extension of Mediterranean-type vegetation in North Africa caused by the increasing mid-to late-Holocene aridification in this region after ca. 7--6 kyr (Hoelzmann & al., 2004; Holmes & Hoelzmann, 2017; Lézine, 2017), which lead to an extension of arid to hyper-arid conditions and a northward spread of Saharo-Sahelian at the cost of Mediterranean flora.

### *Avena breviaristata* -- collected only once

One of the rarest, perhaps even the rarest plant species in Mediterranean North Africa is the grass *A. breviaristata* Barrate, which was collected in 1882 by the French botanist Aristide-Horace Letourneux in Algeria. Letourneux lived between 1881 and 1890 in this country (Stafleu & Cowan, 1979) and was a collaborator of Ernest Cosson (Stafleu & Cowan, 1976) who worked chiefly on the flora of Algeria. Letourneux’ specimen of *A. breviaristata* was incorporated in Cosson’s herbarium. Due to Cosson’s death in 1889, the *Compendium florae atlanticae* remained largely incomplete so that only the first two volumes (general introduction, Ranunculaceae--Cruciferae) were published (Cosson, 1881, 1883--1887). “*Avena breviaristata* Barratte, in litt.” was published as a new species together with a detailed description of characteristics in the monocot volume of the *Flore de l’Algérie* of Battandier & Trabut (1885) and was listed as “*Avena breviaristata* Barratte” in their *Flore analytique et synoptique de l’Algérie et de la Tunisie* (Battandier & Trabut, 1904), but without indicating who had authored the morphological description. Jean François Gustave Barratte was Cosson’s secretary and curator of his herbarium (Stafleu & Cowan, 1976; see also Cosson, 1883--1887 for further information) which was donated to the Muséum National d’Histoire Naturelle in Paris (P) in 1904.

*Avena breviaristata* was subsequently treated in the taxonomic monograph *Contribution à l’étude des Avena sect. Avenastrum (Eurasie et région méditerranéenne)* authored by Alfred Saint-Yves (1931). The author explicitly stated that he had seen the specimen in Cosson’s herbarium (“Spec. exam. -- Algérie: «Ouled Sahari Maio 1882 A. Letourneux» !! herb. Coss.”; Saint-Yves, 1931: 489) and could study it in detail (“[...] que nous l’avons constaté sur l’unique échantillon authentique de l’herbier Cosson et dont nous avons pu faire une minutieuse étude grace a l’extrême bienveillance du Professeur Lecomte à notre égard […]”; Saint-Yves, 1931: 489). Furthermore, Saint-Yves (1931) reported many characters of *A. breviaristata* not mentioned by Battandier & Trabut (1895) and published for the first time drawings of the leaf blade and the awn in transverse sections (Saint-Yves, 1931: figs. 37, 38, pl. IV).

### Type specimen lost some time after 1931

The vol. 2 of the *Flore de l’Afrique du Nord* of René C.J.E. Maire (1953) [French botanist in Algier] that appeared posthumously (eds. M. Guinochet and L. Faurel; see Stafleu & Cowan, 1981) was the next work with a detailed account on *A. breviaristata*. It was based primarily on the character descriptions published by Saint-Yves (1931) and contained a downsized reproduction of Saint-Yves’ (1931) drawings of transverse sections of leaf and awn. Astonishingly, it contained a more comprehensive description of the spikelets than Saint-Yves’ work and the first illustration of a spikelet, which was not explicable so far (see Röser & al., 2009: 32--34), because it was simultaneously stated in the text “Cette plante n’est connue que par un pied unique conservé dans l’Herbier Cosson. Ni nous, ni nos collaborateurs n’avons pu jusqu’ici la retrouver” (Maire, 1953: 308). Later, Quézel & Santa (1962: 120) recorded in their *Nouvelle flore de l’Algérie* vol. 1 on *A. breviaristata* only “un seul exemplaire connu jusqu’ici”.

Later attempts to retrieve the lost specimen of *A. breviaristata* at P by the senior author and by Lange (1995: 29) were unsuccessful. Furthermore, *A. breviaristata* was seemingly never recollected which was especially regrettable in view of the interesting but obviously unclear systematic affiliation of this species.

### Uncertain affinities of *A. breviaristata*

The perennial species *A. breviaristata* was removed from the genus *Avena* L. s.str. and transferred to *Helictotrichon* (Henrard, 1940), *Avenochloa* Holub, nom. illeg. (Holub, 1958) and *Avenula* (Dumort.) Dumort. (Holub, 1962). Due to its comparatively short and straight, non-geniculate awn, *Avena breviaristata* was often accommodated under a separate, monospecific section (section *Brevitrichon* Holub; Holub, 1958, 1962, 1976).

Previously overlooked or wrongly reported morphological characters of *A. jahandiezii* Litard., another North African oat-grass, led to an assignment of this species to a separate subgenus (*H.* subg. *Tricholemma* Röser) and finally genus *Tricholemma* (Röser) Röser (Röser 1989; Röser & al., 2009). This status was meanwhile supported by molecular phylogenetic data (Grebenstein & al., 1998; Wölk & Röser, 2014, 2017). Some morphological characters suggested a relationship between *T. jahandiezii* and *A. breviaristata*. Relying on the descriptions published by the previous authors (i.e., Battandier & Trabut, 1895; Saint-Yves, 1931; Maire, 1953) and, unfortunately, without an opportunity to study the type specimen, the only specimen ever collected, *A. breviaristata* was tentatively transferred to *Tricholemma* by Röser & al. (2009).

### Recovery of the long-lost type specimen and the connected key questions of this study

The recent retrieval of the long-lost type material of *A. breviaristata* and the opportunity to investigate it thoroughly (see Results) now allow us to clarify the classification of this species and related taxa. We hope to answer the following questions: (1) What are the morphological characters of *T. breviaristatum* in detail, especially the yet incompletely studied floral structures (e.g., spikelet parts such as lemma, lodicules)? (2) Does *T. breviaristatum* share derived (synapomorphic) characters with *T. jahandiezii*, which would taxonomically corroborate their inclusion in the same genus, i.e. *Tricholemma*? (3) Which characters can be used to distinguish morphologically superficially similar genera of grasses, especially *Avenula* s.str. *Helictochloa* and *Helictotrichon*. (4) Using this information on diagnostic characters we finally re-evaluate the presumed intergeneric hybrid origin (Soreng & Davis, 2000: 71; Gillespie & al., 2008; Kellogg, 2015) of widespread temperate Eurasian *Avena pubescens* Huds., the nomenclatural type and sole species of the genus *Avenula* as currently circumscribed (Romero Zarco, 2011).

## MATERIAL AND METHODS

### Plant material

The following plant material was used for microscopic examination and microphotography: (1) The type specimen of *Avena breviaristata*: [Algeria, Hauts Plateaux], Ouled Sahari, Maio 1882, A. Letourneux *s.n.* (P00152040); (2) *Tricholemma jahandiezii* (Litard.) Röser, nomenclatural type (Röser, 1989: 46) of genus *Tricholemma*: Morocco, Moyen Atlas, Ifrane, 4 Oct 1995, M. Röser 10297, pot plants grown in the Botanical Garden Halle, 10 Mar 2005 (HAL); (3) *H. sempervirens* (Vill.) Pilg., nomenclatural type (Schweickerdt, 1937) of genus *Helictotrichon* encompassing in total ca. 18 species (Wölk & Röser, 2017): Germany, Saxony-Anhalt, Blankenburg, planted on a traffic island, 14 May 2016, M. Röser 11226 & N. Tkach (HAL); (4) *Helictochloa pratensis* (L.) Romero Zarco, member the genus *Helictochloa* Romero Zarco encompassing in total ca. 24 species (Romero Zarco, 2011): Germany, Saxony-Anhalt, Thale, 14 May 2016, M. Röser 11228 & N. Tkach (HAL); (5) *Avenula pubescens* (Huds.) Dumort., nomenclatural type (Chase, 1939: 568) of the monospecific genus *Avenula*; Germany, Saxony-Anhalt, Warnstedt, 14 May 2016, M. Röser 11231 & N. Tkach (HAL). The plant material consisted of herbarium specimens for species 1--5 and fresh inflorescences preserved in 70% ethanol for species 3--5.

### Trait selection and microscopic study

For a comparison of the species in question several morphological characters were evaluated: (1) Leaf blades were studied in regard to the type of the enfolding by the presence of parallel longitudinal adaxial furrows and ribs or by only two rows of bulliform cells parallel to a distinctive midrib. Additionally, the abaxial structure of the sheaths of basal leaves was examined. (2) On the transition between leaf sheath and leaf blade the differently structured ligules are located. These structures are formed by the adaxial epidermal cells and enclose the stem in flowering or the next inner leaf in sterile shoots. Ligules are either entirely membranous or have an apical fringe of hairs. (3) The total length of the spikelet was measured from the base of the lower glume to the apex of the uppermost lemma or to the apex of the upper glume if it exceeds the lemmas. In addition, the uppermost floret was recorded as fertile or reduced. (4) The hair length of the callus was measured and the position of the callus recorded. The disarticulation of the spikelet axis (rachilla), either below each callus (floret) or only below the callus of the lowermost lemma (floret), determines the fruit dispersal by being the breaking point of those components of the spikelet that serve as diaspores. (5) The length of the rachilla segment between the lowermost and the second lemma was measured. The segments were examined for their hairiness and it was distinguished between lower and upper part if necessary. (6) The number, total length and form, as well as length of nerves or keels, existence of apical notches and hairiness were recorded for glumes, lemmas and paleas. Additionally, the type of hairs was determined in either ad- or abaxial position of the rachilla according to Ellis (1979) as normal or as apically pointed and basally thickened prickle hairs. (7) Awns of the lemma were measured for length, hairiness and position of the bend, if present. The latter is formed at the transition between a longitudinally twisted basal part (columna) of the geniculate awn and a straight upper part (subula). (8) The lodicules were examined as excellent trait in morphological comparison of the taxa in question due to their genus- and sometimes species-specific shape (Röser, 1989; Lange, 1995; Romero Zarco, 2011).

### Methods and data analysis

In accordance to the selected morphological criteria, all plant parts were prepared and measured in dry or wet state under a Zeiss Stemi 1000 incident light microscope. The structure of lodicules was studied in bright and dark field on a Zeiss Axioskop 2 transmitted light microscope. Photographs were taken using a computer-assisted camera (Axiocam) and Zeiss Axiovision software for measurements. Supplementary drawings were made for ligules and three-dimensional lemma structure. As for the investigation, spikelets of an average size were selected to minimize errors by measurements of extreme values in upward or downward ranges. Subsequently, all data and measured values were tabulated.

## RESULTS AND DISCUSSION

### Recovery of the holotype of *Avena breviaristata* at P

In 2012 while checking some new online digital images at P (https://science.mnhn.fr/all/search) MR found a database entry for *A. breviaristata.* In addition, an excellent scan of the long-lost type specimen of *A. breviaristata* from Cosson’s herbarium was retrieved (see http://coldb.mnhn.fr/catalognumber/mnhn/p/p00152040), also found in *JSTOR Global Plants* (https://plants.jstor.org/collection/TYPSPE) and reproduced in the Electr. Suppl.: Fig. S3]. The sheet (P00152040) contains one flowering shoot with two additional vegetative, sterile shoots and a separate non-flowering vegetative shoot, all of which have their basal leaves and leaf sheaths (Electr. Suppl.: Figs. S1, S3). The label placed at the bottom in the centre is the original label written by Letourneux: “*Avena* [,] Ouled Sahari [,] Maio 1882 [,] A. Letourneux” (Electr. Suppl.: Figs. S1, S2A, S3). According to the protologue (Battandier & Trabut, 1895) these shoots, collected in the field probably from to same plant specimen, represent the holotype (Turland & al., 2018: Art. 9.1, Note 1 of the ICN 2018). The herbarium sheet as image available online on the web pages of P and in *JSTOR Global Plants* (Electr. Suppl.: Fig. S3) has an additional label, which was carefully bent back in our Electr. Suppl.: Fig. S1. It bears the handwritten name “*Avena breviaristata* Barratte”, on which a smaller label “TYPE” and a barcode label are attached. Moreover, the herbarium sheet includes the hand-drawn original illustration of the leaf blade in transverse section made by Saint-Yves that served as printing template for the illustration in his monograph of 1931 (see above). A leaf fragment is attached to this drawing (Electr. Suppl.: Fig. 2B). Finally, at the top of the sheet Barratte’s handwritten transcript of a letter he had sent to Trabut is attached (Electr. Suppl.: Fig. S5). This transcript was temporarily removed for making our image of this plant specimen in Electr. Suppl.: Fig. S1 and for the online images available on the web pages of P and in *JSTOR Global Plants* cited above.

### Barratte’s original letter to Trabut and a fragment of the holotype at MPU

To our great surprise, Barratte’s original handwritten letter sent to Trabut is still preserved in the collection of R. Maire at MPU. Images of the letter are available online under https://science.mnhn.fr/institution/um/collection/mpu/item/mpu001465. Due to its considerable importance and the historical information on the type collection it is reproduced as Electr. Suppl.: Fig. S4. For our transcript of the letter see Electr. Suppl.: Appendix S1.

Our English translation of this letter reads as follows: “Paris, 19 January 1894. Sir and dear collegue, my first idea was to send you the *Avena* collected in May 1882 by Letourneux in the Ouled Sahari, but the only flowering specimen that exists in the herbarium [i.e., the herbarium of Cosson] is folded in a manner that does not comply with the postal requirement; I could send it as a parcel but I believe it is better to give you a description than to damage it during the transport. In addition to these few lines I enclose one of the better spikelets based on which you can verify and complete the characters. Here is the description of this nice species made on the single specimen Letourneux collected, which is unfortunately devoid of sterile leaf shoots; you may delete what seems useless to you. [At this point follows the description of the morphological characters, widely corresponding with that published by Battandier & Trabut (1895) except for some details (see below)] This description is undoubtedly a little bit long but I repeat that you can make omissions as you consider it necessary. Furthermore, I tell you that Letourneux wrote nothing else on his label than «Ouled Sahari». Are you sure that this is on Dj. Senalba, where he has collected this nice species? I went through the letters written to Mr. Cosson during that year [i.e., 1882] but I found nothing about this. I would perhaps be happier, if I had the time to look at the correspondence of the following years; I will do this as soon as I have an opportunity to do so. The species name *Senalbensis* has certainly not yet been given to a species of *Avena* but wouldn’t it be better to give this nice species a name that matches one of its main characters, *breviseta* or even better *breviaristata*? Sir and dear colleague, sincerely yours, G. Barratte.”

This letter at MPU (Electr. Suppl.: Fig. S4; sheet MPU001465a) is literally the same as Barratte’s transcript at P, which bears angled in the upper left corner the words “Copie de la lettre addressée au Dr. Trabut” handwritten by Barratte (Electr. Suppl.: Fig. S5). Interestingly, the letter at P (“copie”) was actually Barratte’s draft, which is evident from several deletions and insertions. Thereafter he sent a clear copy (now at MPU), besides that made in more decent handwriting, to Trabut and kept the draft for his own records.

Barratte’s description was widely adopted in the protologue of *A. breviaristata* published by Battandier & Trabut (1895), only the statement “Plante […] rameuses dans leur partie supérieure […].” was omitted in this work, which is appropriate, because the culm is not branched but only the inflorescence. However, the inflorescence is comparatively expansive, richly branched and the branches and pedicels are rather long. The description of the spikelets and the spikelet parts was modified and extended, which seems to be due to Trabut’s study of the spikelet that Barratte included by with his letter to Trabut as he wrote (see above and our transcription of the letter below. However, the disarticulation of the spikelet only above the lowermost lemma remained unnoticed in this species, until it was recognized by Saint-Yves (1931: 488): “flos inferior solus articulatus”. Second, Saint-Yves (1931: 489, “Trabut n’a pas vu la plante”) correctly argued that Trabut never had seen the whole plant of Cosson’s herbarium, because then he would have noticed that the ligules are not short and glabrous as written by Barratte and accepted in the protologue. Trabut only knew the spikelet and not the whole plant specimen that was not sent to him (see Barratte’s letter).

Astonishingly, this spikelet is still preserved in Maire’s herbarium at MPU (ditto https://science.mnhn.fr/institution/um/collection/mpu/item/mpu001465; also https://collections.umontpellier.fr/collections/botanique/herbier-mpu and https://plants.jstor.org/collection/TYPSPE; reproduced in Electr. Suppl.: Fig. S6). Altogether, there are two spikelets with their pedicels present on sheet MPU001465 (Electr. Suppl.: Fig. S2C), one of which disintegrated into the pedicel with the lower and upper glume and the compound structure of all florets, due to the disarticulation of the floret axis only above the upper glume. This spikelet was most obviously the template for the previously enigmatic first illustration of a spikelet of *A. breviaristata* in Maire’s *Flore de l’Afrique du Nord* (1953), because the type specimen was already lost at that time (see above). The presence of a second spikelet is not explicable from Barratte’s letter such as the additional presence of a leaf fragment (Electr. Suppl.: Fig. S2C), which resembles that used by Saint-Yves on voucher P00152040 for sectioning the leaf (Electr. Suppl.: Fig. S2B). The sparse plant material at MPU, however, must be regarded as fragment removed from, and belonging to the holotype and not as a duplicate or something like that, which would represent an isotype. It is correctly labelled at the bottom of the sheet MPU001465 as “fragmentum typi”.

### Locus typi

Barratte’s letter from 1894 stated that Letourneux only wrote “Ouled Sahari” as type locality on the original label and questioned that this agrees with “Dj. [Djebel] Senalba”, a location seemingly invoked already before by Trabut. The protologue to *A. breviaristata* finally states “Ouled-Sahari entre Boghari et Bou Saâda au-dessus du Zahrès-Chergui” (Battandier & Trabut, 1895: 184), which was largely reiterated by Maire (1953: 358) and Quezel & Santa (1962: 120). The locality “Ouled Sahari” and the latter three publications thus probably refer to a town now named Had-Sahary (N 35°21’02” E 3°21’55”, 840 m a.s.l.), which is located ca. 20 km NW Zahrez Chergui. Ouled Sahari is not listed among the botanical collecting sites in Algeria by Cosson (1881: 262; see entry no. 70 on Djelfa and surroundings) but is contained in an updated list published in the following year (Cosson, 1882: 140; see also entry no. 70), possibly due to the recent collection of Letourneux.

### Morphology

Our survey of morphological characters based on recent investigation (see Material and methods) and data from the literature shows that *Tricholemma breviaristatum* shares several characters of vegetative morphology with other genera (Table 1). The leaf anatomy and folding pattern of *T. breviaristatum* basically agrees with *Avenula*, *Helictochloa* and *T. jahandiezii* rather than with *Helictotrichon*. The same applies to the ligules of the leaves, which are membranous in *T. breviaristatum* and *T. jahandiezii* (Fig. 1A, B), a rare characteristic in *Helictotrichon* but the rule in *Avenula* and *Helictochloa* (Table 1).

**Table 1.** Comparison of leaf and spikelet characters between *Tricholemma breviaristatum* and morphologically similar taxa traditionally unified under the broadly delineated genera *Helictotrichon* and *Avenula*. Data based on own observations or taken from the literature (mainly Saint-Yves, 1931; Tzvelev, 1976; Lange, 1995; Wu & Phillips, 2006).

|  | *Tricholemma breviaristatum* | *Tricholemma jahandiezii* | *Helictotrichon* s.str. <sup>a)</sup><br>(ca. 20 species) | *Helictochloa*<br>(ca. 24 species) | *Avenula* s.str.<br>(1 species) |
| --- | --- | --- | --- | --- | --- |
| Upper surface of leaf blade | with 2 rows of bulliform cells | with 2 rows of bulliform cells | longitudinally ribbed, with several deep furrows | with 2 rows of bulliform cells | with 2 rows of bulliform cells |
| Folding of leaf blade | conduplicate | conduplicate | convolute | conduplicate | conduplicate |
| Ligule | membranous | membranous | fringe of hairs, rarely membranous | membranous | membranous |
| Disarticulation of the spikelet | only above glumes | only above glumes | above glumes and between the florets, rarely only above glumes <sup>b)</sup> | above glumes and between the florets | above glumes and between the florets |
| Lemma back with prominent midvein | yes | yes | no | no | no |
| Midvein of lemma back with a row of long hairs | yes | yes | no | no | no |
| Back of lemma in the upper 2/3 with long, spreading macrohairs | yes | yes | no | no | no |
| Structure of awn | only subula | columna and subula | columna and subula | columna and subula | columna and subula |
| Shape of columna | – | almost cylindrical | almost cylindrical | flattened | almost cylindrical |
| Palea keels | ciliolate | ciliolate | ciliolate | ciliolate | glabrous |
| Lodicules | ovate-lanceolate, margin smooth, >2 mm long | ovate-lanceolate, margin smooth, >2 mm long | narrowly triangular, margin smooth, >>2 mm long | ovate-lanceolate, with lateral lobe, margin smooth, >>2 mm long | broadly ovate-lanceolate, margin and apex notched, <1 mm long |

<sup>a)</sup> Excluding *Trisetopsis* Röser & A.Wölk, *Tzveleviochloa* Röser & A.Wölk and *×Trisetopsotrichon* Röser & A.Wölk. <sup>b)</sup> In a group of species endemic to the Alps (*H. parlatorei* (J.Woods) Pilg., *H. setaceum* (Vill.) Henrard, *H. sempervirens*) and the Atlantic, west Mediterranean species *H. thorei* Röser and *H. pallens* (Link) J.M.Couderc & Guédès. The latter two species are sometimes accommodated under the separate genus *Pseudarrhenatherum* Rouy.

**Fig. 1.**
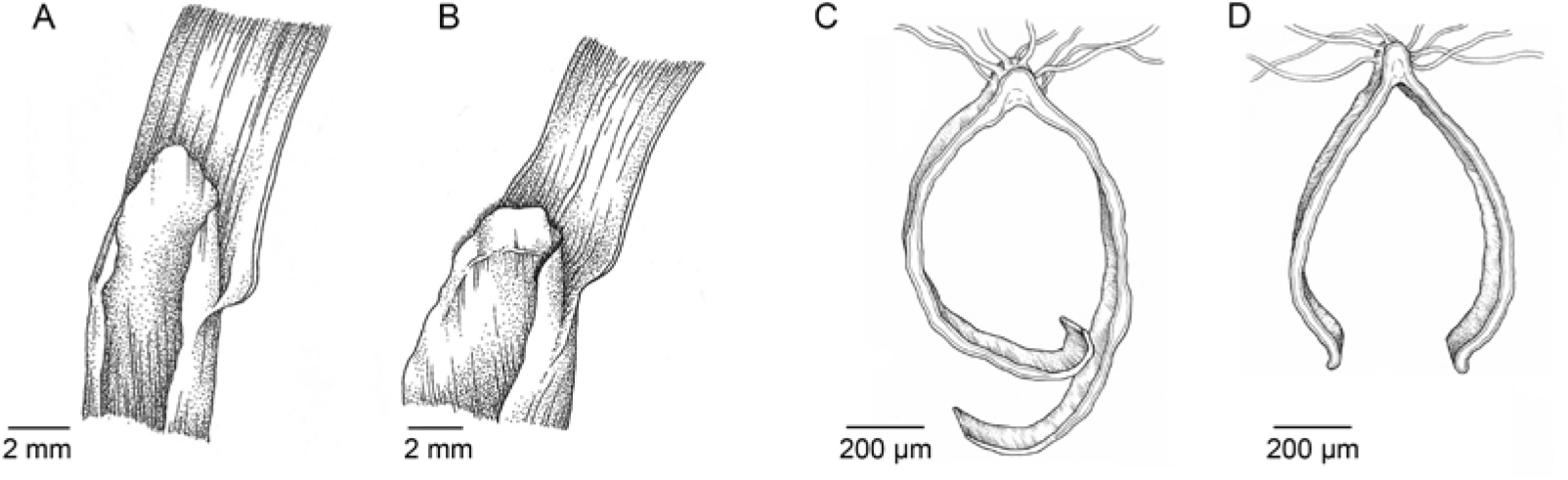
**A--B**, Ligules of upper cauline leaves: **A**, *Tricholemma breviaristatum*; **B**, *T. jahandiezii*. Note the hairs on the upper margin of the ligule. **C--D**, Lemmas in transverse section below the insertion of the awn: **C**, *Tricholemma breviaristatum*; **D**, *T. jahandiezii*. Note the prominent midvein of the lemma back and the row of stiff macrohairs.

A disarticulation of the spikelets only above the glumes is shared between *T. breviaristatum* and *T. jahandiezii*. It should be noted that the disarticulation of the spikelet was erroneously reported for the latter species as below each floret (Saint-Yves, 1931: 424: “flores omnes articulati et decidui” and Maire, 1953: 298: “Fleurs toutes articulées sur la rachéole […]”). This shared type of spikelet disintegration only above the glumes, resulting in synaptospermy, is absent in the other taxa in question except for certain species of *Helictotrichon* (Table 1, footnote b). The majority of species of *Helictotrichon* shares a disarticulation of the rachilla below each floret with *Avenula* and *Helictochloa*, representing the evolutionary plesiomorphic state. *Tricholemma breviaristatum* and *T. jahandiezii* share uncommon characters of the lemma. The lemmas have raised midveins below the insertion of the awn. Along this prominent midvein the lemma additionally bears a conspicuous line of long, white macrohairs. Both characters are missing in the other taxa. They are a symplesiomorphy for *T. breviaristatum* and *T. jahandiezii*. The back of the lemma additionally has in the upper two thirds a more (*T. jahandiezii*) or less (*T. breviaristatum*) dense indumentum of spreading, long macrohairs (Fig. 2 A, B), which is absent in the other taxa. A unique feature of *T. breviaristatum* is the bristle-like shape of the awn, which is straight and not twisted in the lower part as in the other taxa (Figs. 2A, B, 3A; Table 1). The awn has become reduced to the subula. It may be argued that this reduction process could partly result from the mode of dispersal (synaptospermy) caused by the disarticulation of the spikelets only above the glumes. Such species, however, tend to have downsized or lost awns in the lemmas of the upper florets as noted for *Helictotrichon sempervirens* (Röser 1989: 75), the species previously accommodated under *Pseudarrhenatherum* Rouy [*H. pallens* (Link) J.M.Couderc & Guédès, *H. thorei* Röser; Romero Zarco, 1985, 2011] or the genus *Arrhenatherum* P.Beauv. (Clayton & Renvoize, 1986; Watson & Dallwitz, 1992). The correlations however between the type of spikelet disarticulation and the development of awns, are not absolutely firm, because the lowermost floret (lemma) has a well-developed awn in all above-mentioned taxa, in contrast to *T. breviaristatum*. However, *T. jahandiezii* has a well-developed awn not only in the lower, but also in the upper lemmas of the spikelets despite sharing the disarticulation of the spikelets only above the glumes. The unique awn structure in *T. breviaristatum* is an autapomorphic character of this species such as the flattened column of the awn in the species of the genus *Helictochloa*. The palea of *Avenula* (only *A. pubescens*) is unique in having glabrous keels, an autapomorphic character of this genus. The shape and size of lodicules of *T. breviaristatum* and *T. jahandiezii* are rather similar and differ from that of *Helictochloa* and *Helictotrichon*. *Avenula* deviates strongly from all of them due to its very short and deviantly shaped lodicules with a side lobe (Table 1, Fig. 4).

**Fig. 2.**
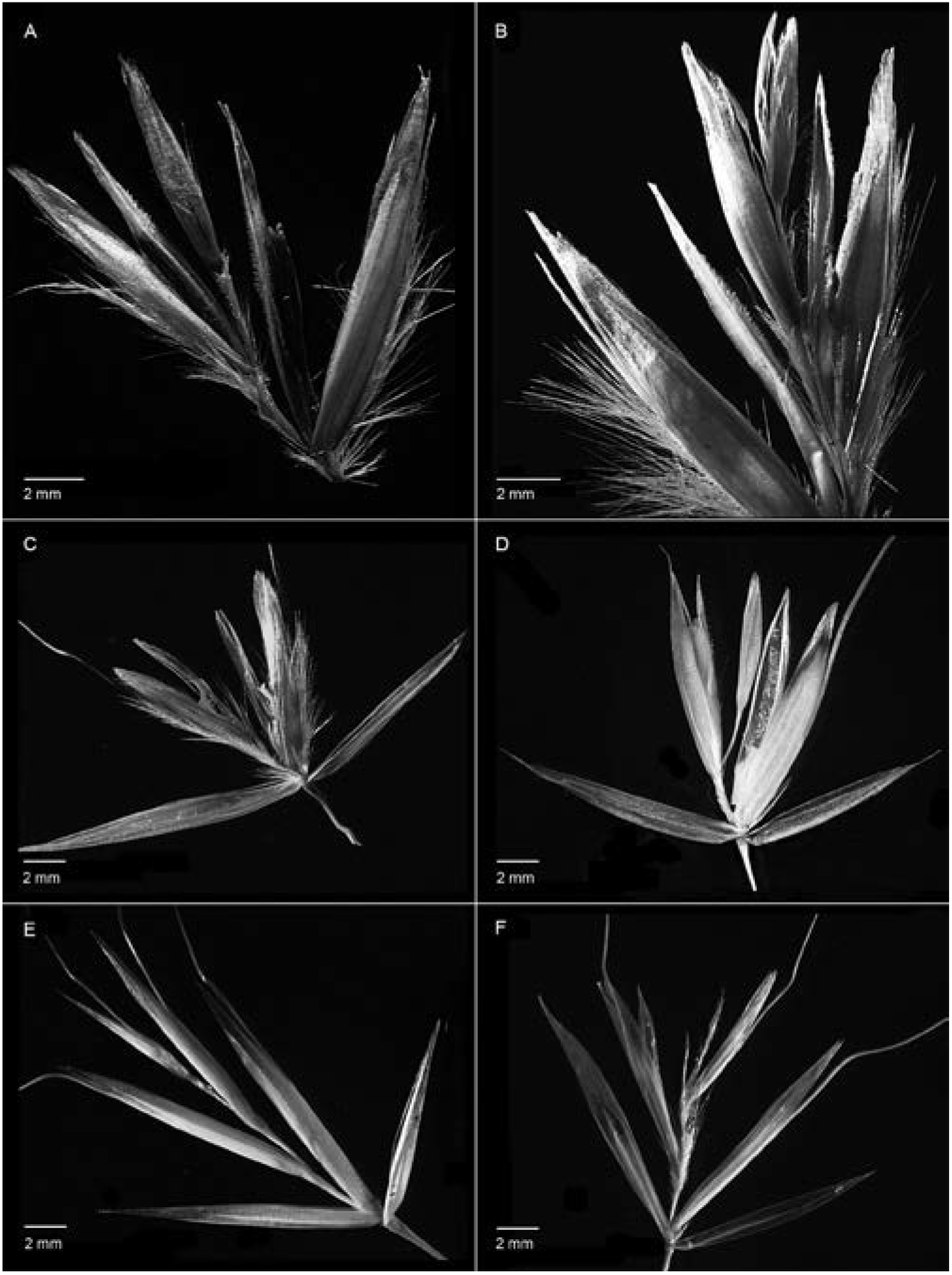
Spikelets in lateral view (dark field): **A--B**, *Tricholemma breviaristatum* (without lower and upper glume); **C**, *T. jahandiezii*; **D**, *Helictotrichon sempervirens*; **E**, *Helictochloa pratensis*; **F**, *Avenula pubescens*.

**Fig. 3.**
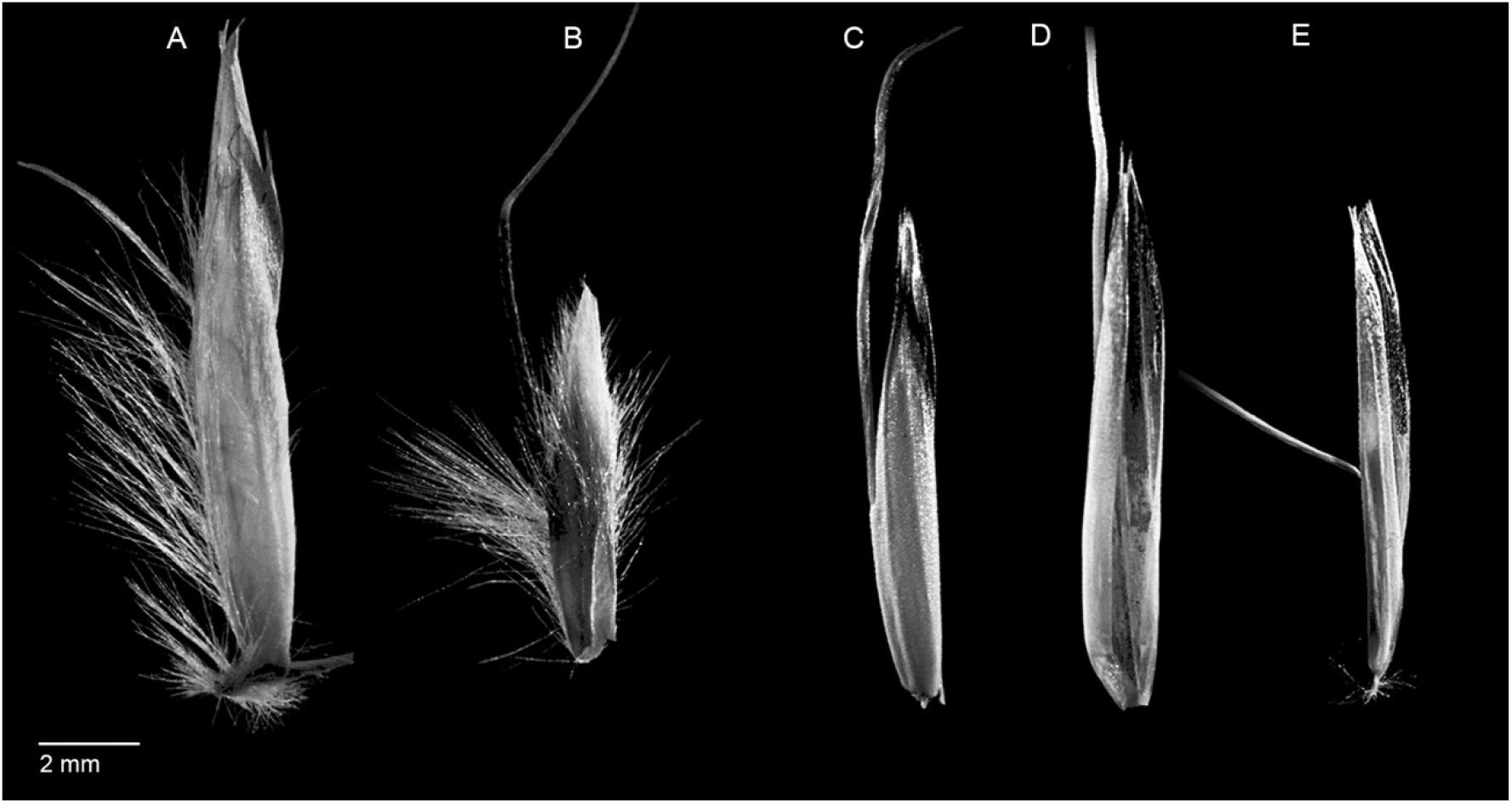
Lemmas in lateral view with insertion of the awn (dark field): **A**, *Tricholemma breviaristatum;* **B**, *T. jahandiezii*; **C**, *Helictotrichon sempervirens*; **D**, *Helictochloa pratensis*; **E**, *Avenula pubescens*.

**Fig. 4.**
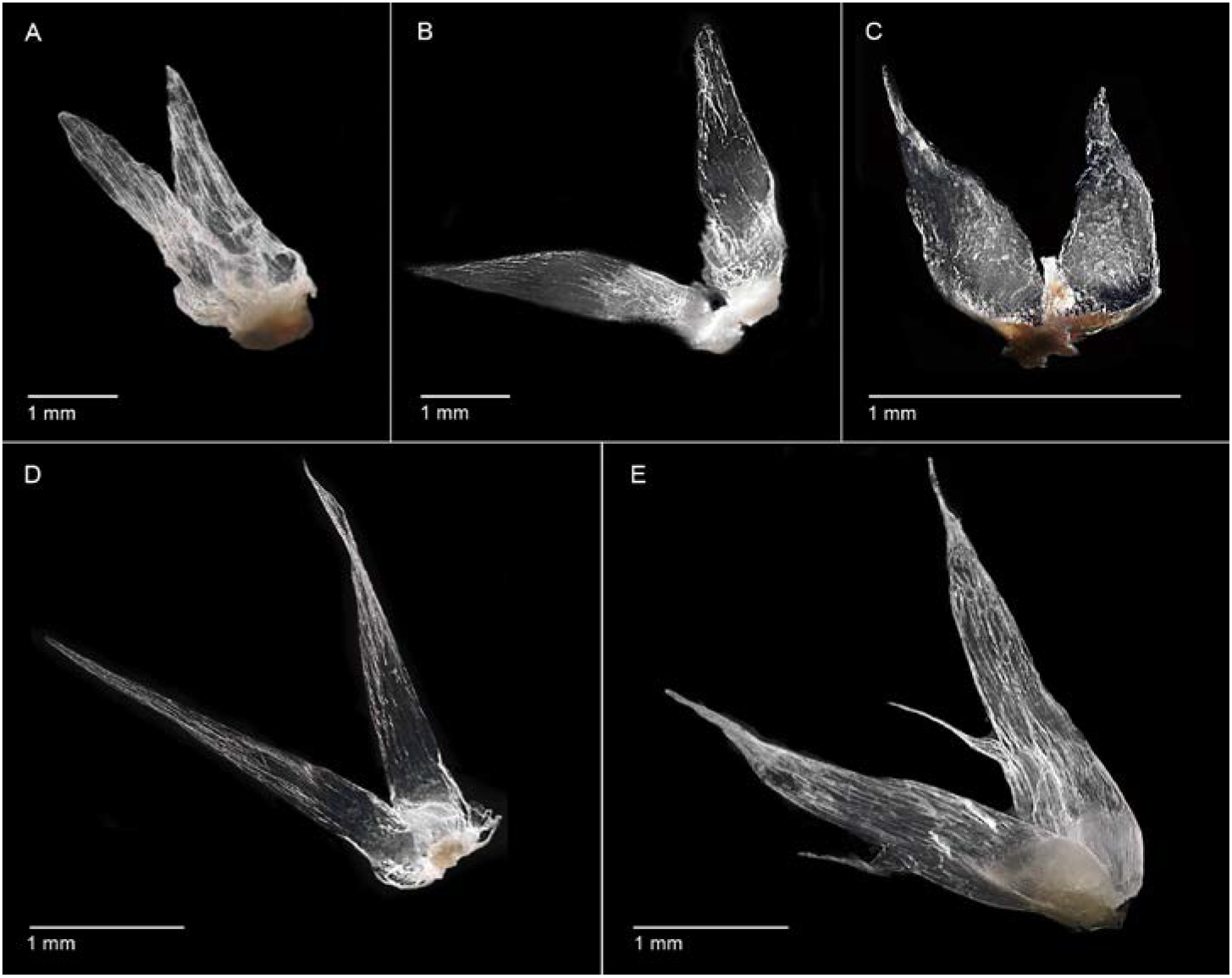
Lodicules (dark field): **A**, *Tricholemma breviaristatum*; **B**, *T. jahandiezii*; **C**, *Avenula pubescens*; **D**, *Helictotrichon sempervirens*; **E**, *Helictochloa pratensis*.

## CONCLUSIONS

The morphological characters surveyed in the rediscovered type specimen of *Avena breviaristata* suggests a close relationship to *T. jahandiezii*, especially due to uncommon features of the lemma (prominent midrib with a row of hairs), the same and comparatively rare type of rachilla disarticulation and similarly shaped lodicules. This supports the inclusion of both species in the genus *Tricholemma* (Röser & al., 2009). This is especially striking in the overall appearance of the inflorescences and the spikelets with long glumes (Fig. 5A, C). Although being larger in *T. breviaristatum* than in *T. jahandiezii* the inflorescence is comparatively expansive and richly branched in both species. Moreover, the branches are rather long, slender and flexuous, smooth and without prickle hairs. The pedicels are almost unthickened below the glumes, which is uncommon among the taxa studied. Uncommon is also the dense indumentum of the basal leaf sheaths of both species with reflexed hairs directed downwards (Fig. 5B, D).

**Fig. 5.**
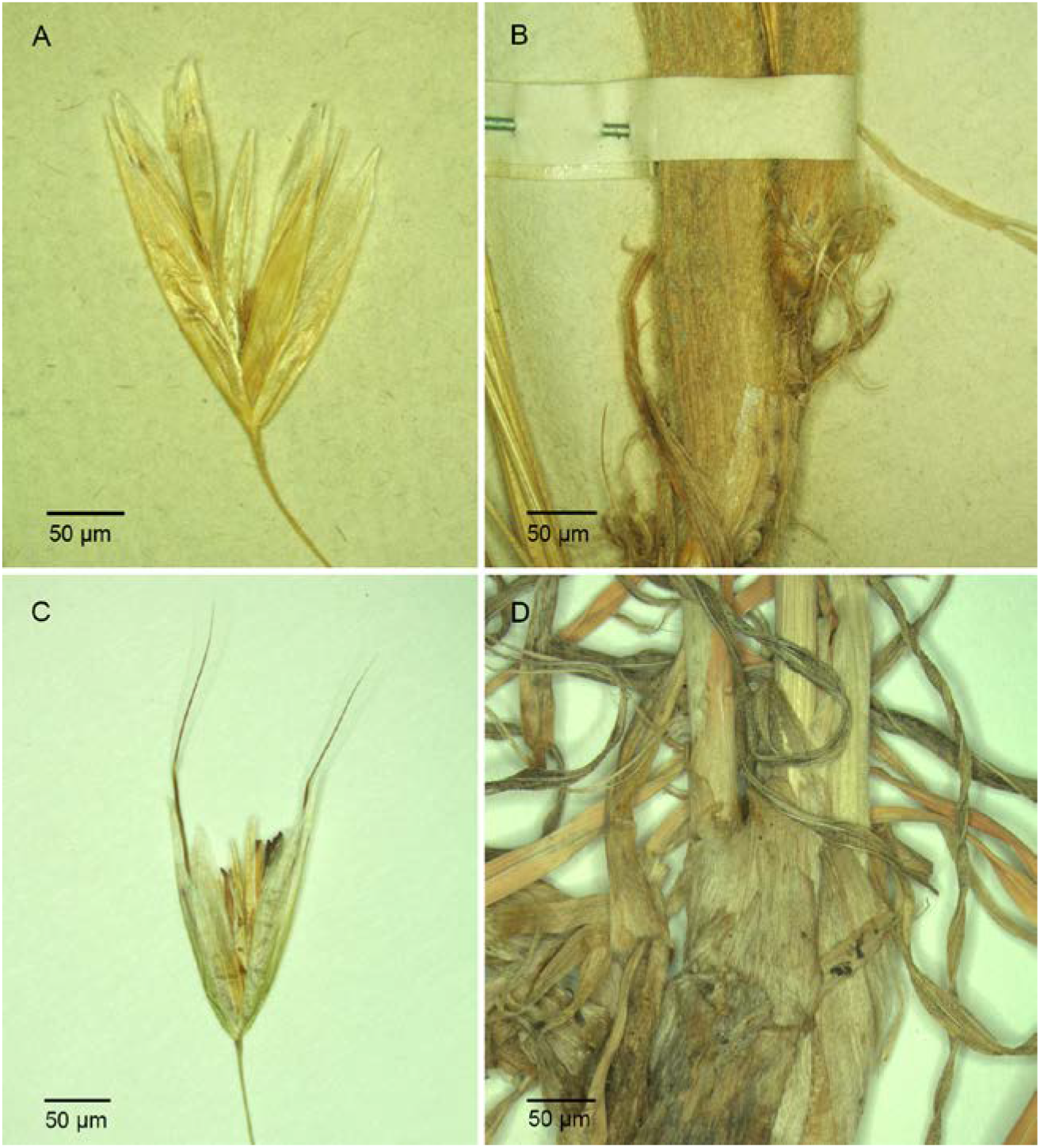
Spikelets (**A**, **C**) and basal leaf sheaths (**B**, **D**): **A--B**, *Tricholemma breviaristatum*; **C--D**, *T. jahandiezii*.

Nevertheless, *T. breviaristatum* stands out by its unique awn structure that is best interpreted as reduction, in some way probably induced by its disarticulation of the floret axis, which is shared with *T. jahandiezii*. The habitat of *T. breviaristatum*, in the Algerian Hauts Plateaux in an altitude of ca. 840 m probably under comparatively arid conditions and in open vegetation, may play a role in the dispersal of diaspores, for which possession of an elaborate awn is eventually disadvantageous. The situation for diaspore dispersal in *T. jahandiezii*, distributed in montane altitudes in the comparatively humid Moyen Atlas is likely quite different (Maire, 1953; Röser, 1989; Ibn Tattou, 2014). Elaborate awns with a column capable of hygroscopic movement might be a selection advantage in this region.

A strongly different climatic adaptation of both species is evident also from their leaf architecture, which is highly xeromorphic in *T. breviaristatum* (junciform leaf, subepidermal strands of sclerenchyma, abaxial furrows with embedded stomata, etc.) but mesomorphic in *T. jahandiezii* with comparatively thin, flat leaves and no sclerenchyma (Röser, 1989, 1996). The occurrence of a xeromorphic *versus* mesomorphic, climatically differently adapted species pair in the relic, disjunctively distributed North African genus *Tricholemma* (Fig. 6) resembles the situation in Mediterranean members of the much more widespread genus *Helictochloa* as currently understood, in which xeromorphic, partly highly polyploid taxa originated in different species complexes from diploid and primarily mesomorphic ancestors (Röser, 1996). To address this cytogenetic question and to study the dispersal and ecological behavior of *T. breviaristatum* a rediscovery of this species in the field would be essential.

**Fig. 6.**
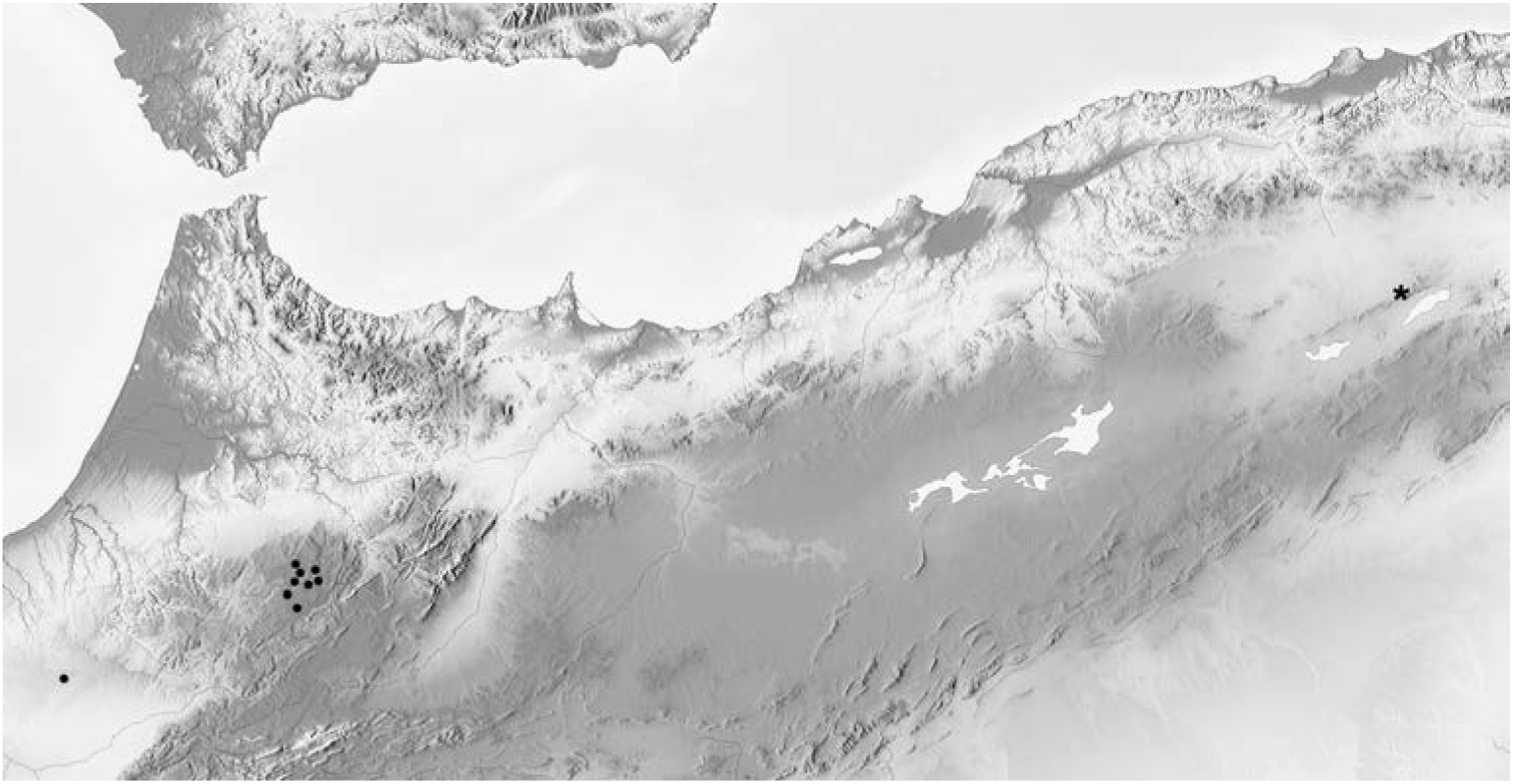
Occurrences of the disjunctively distributed endemic North African genus *Tricholemma*: Asterisk, *T. breviaristatum*; dots: *T. jahandiezii*.

Considering the suggested hybrid origin of *Avenula* (*A. pubescens*) our results make it unlikely that this holds true, at least none of the taxa examined here appears as potential parent candidate. Especially the very odd character of glabrous lemma keels and the peculiar shape of the lodicules are not ‘intermediate’ between any of the other genera studied.

## ACKNOWLEDGEMENTS

We are very grateful to T. Haevermans, M. Jeanson and M. Narfin (P) for sending us the type specimen of *Avena breviaristata* on loan.

---

**Electr. Suppl.: Fig. S1.**
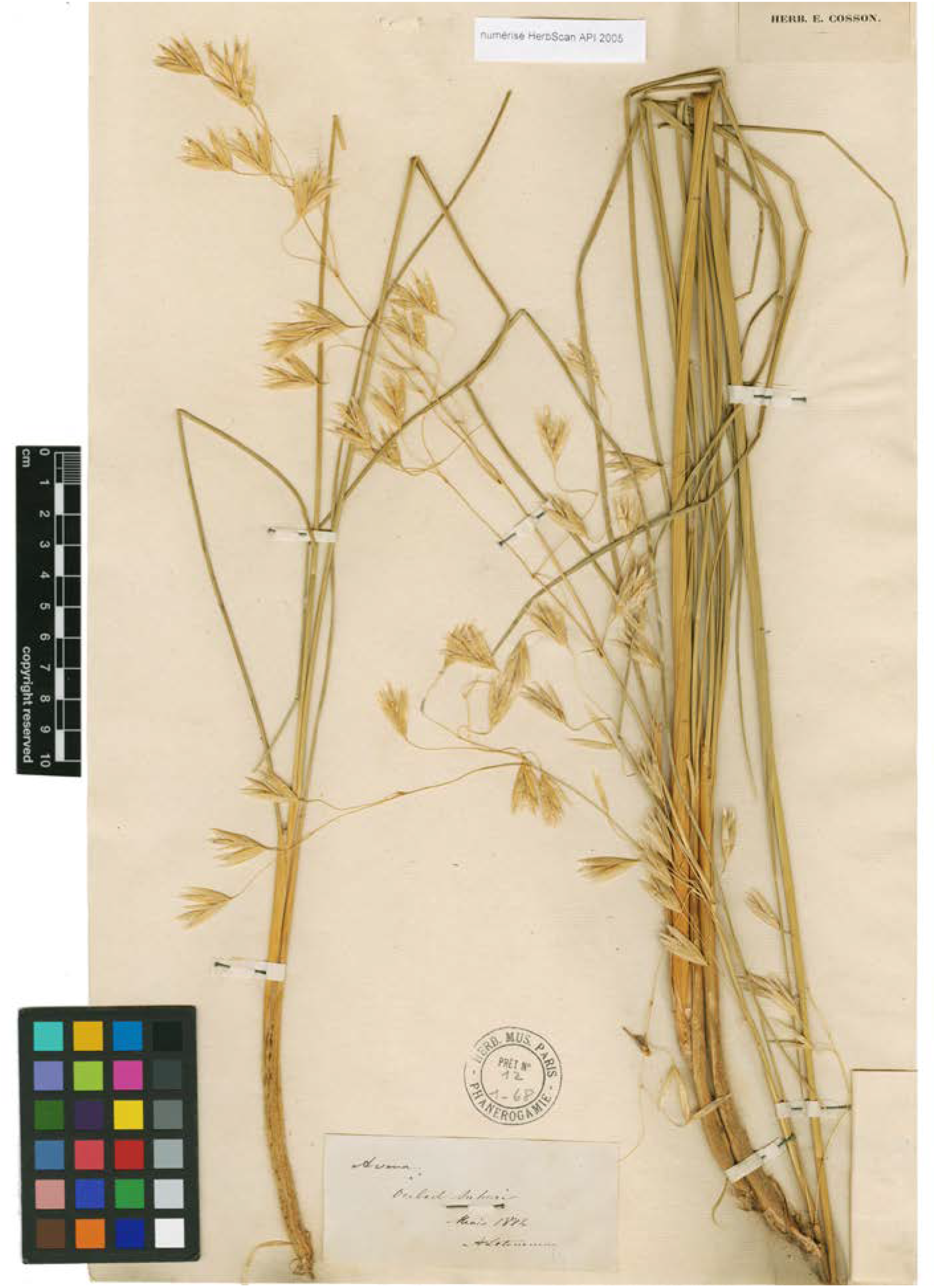
Holotype of *Avena breviaristata*, herbarium sheet P00152040 of the Muséum national d’Histoire naturelle in Paris with the original label of the collector A. Letourneux. Newly made image showing the basal parts of both shoots, which were probably collected in the field from the same plant. A label in the lower right hand corner of the sheet bearing an attached barcode label was carefully bent back (see Electr. Suppl.: Fig. S3). A handwritten transcript of a letter of Barratte to Trabut attached by a pin was temporarily removed for the image (see Electr. Suppl.: Fig. S5). A small printed label not mounted on the sheet (“numérisée HerbScan API 2005”) indicates that this sheet was scanned and perhaps actually recovered in the course of the *African Plant Initiative*, an international project on type specimens of African plant species launched by the Andrew W. Mellon Foundation, U.S.A. This label was hidden behind a drawing of the leaf blade in transverse section made by Saint-Yves (Electr. Suppl.: Fig. S2B), which was not mounted on the sheet (see text) and temporarily removed for this image.

**Electr. Suppl.: Fig. S2.**
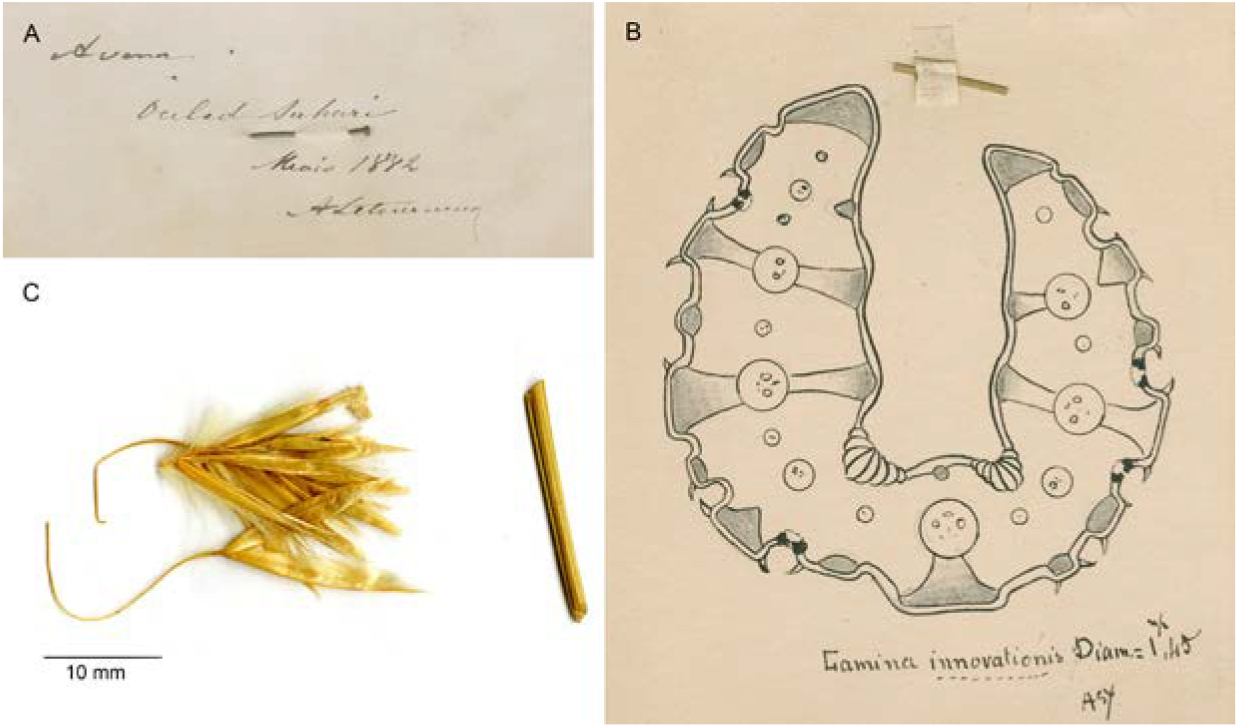
Details of the herbarium sheets of *Avena breviaristata* in P and MPU. **A**, Original label written by A. Letourneux on P00152040. **B**, Drawing of the leaf blade in transverse section made by Saint-Yves, which served as printing template in his monograph, placed on the sheet P00152040. **C**, Detail from sheet MPU001465 with one intact spikelet and overlying a second one disarticulated above the upper glume. The latter spikelet seemingly was the template for the otherwise inexplicable first illustration of a spikelet of *A. breviaristata* in Maire (1953). See text for further explanation.

**Electr. Suppl.: Fig. S3.**
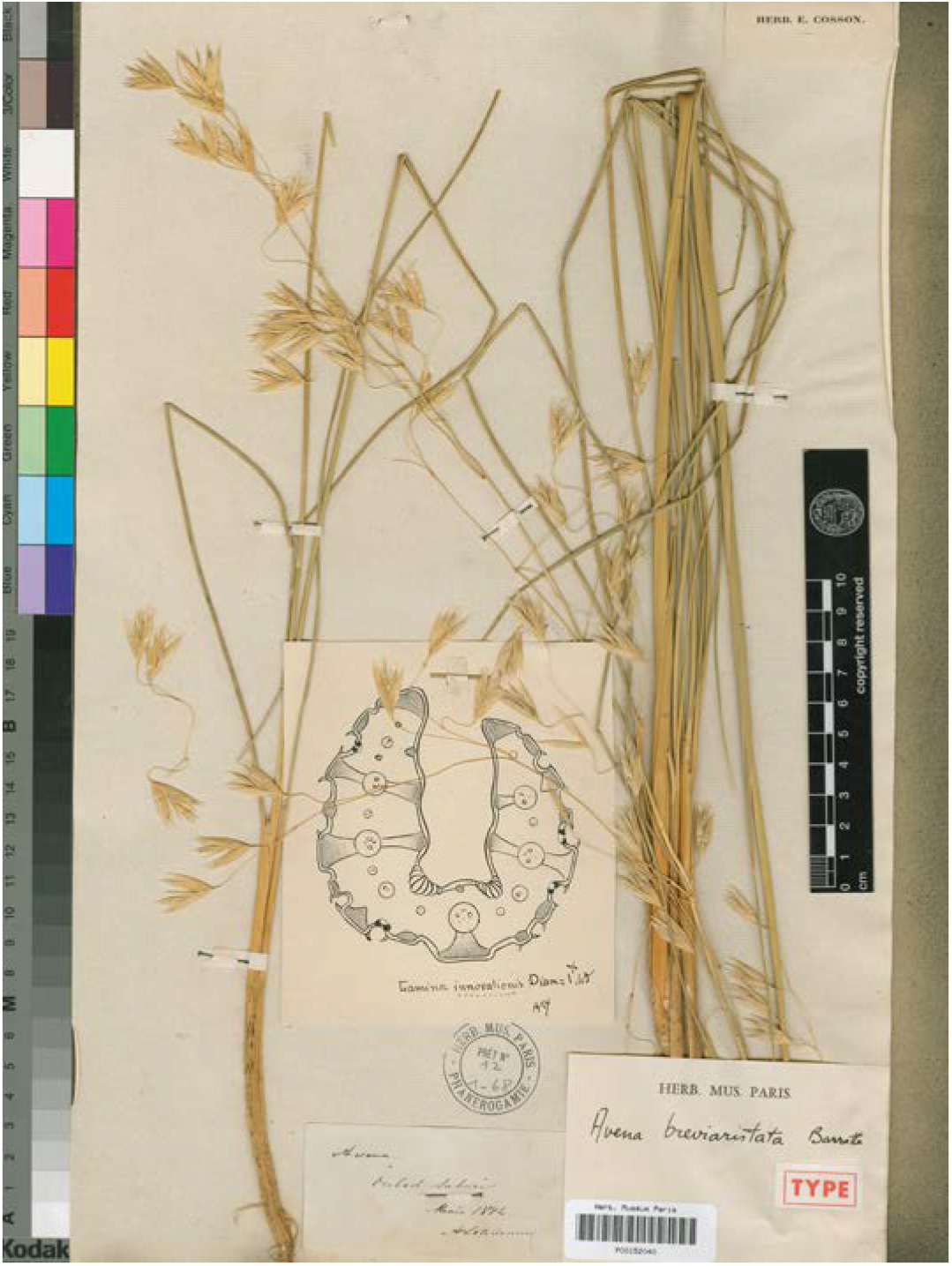
Type specimen (holotype) of *Avena breviaristata* (P00152040) available under http://coldb.mnhn.fr/catalognumber/mnhn/p/p00152040 and https://plants.jstor.org/collection/TYPSPE.

**Electr. Suppl.: Fig. S4.**
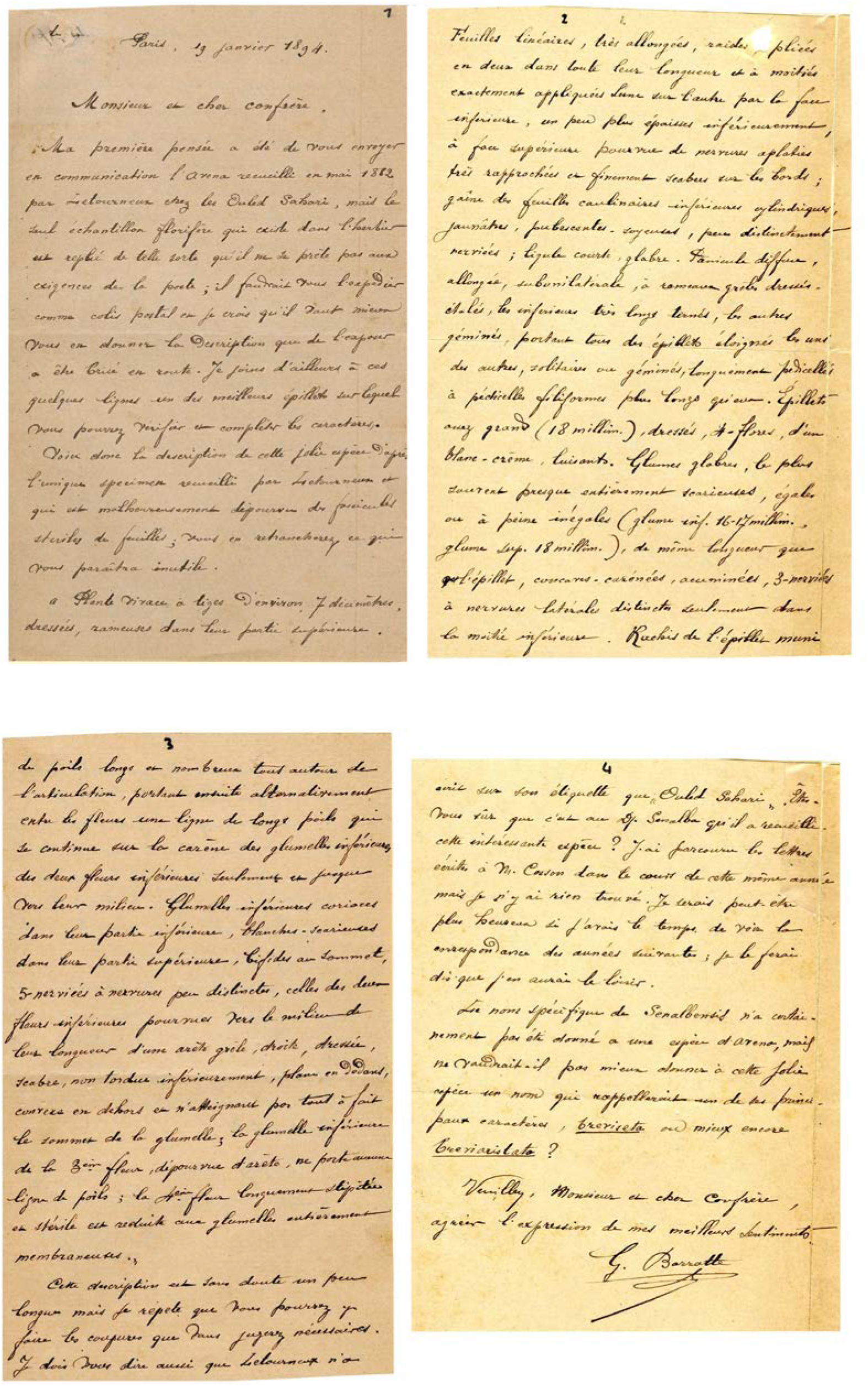
Handwritten original letter of Barratte sent to Trabut in 1884, preserved in the collection of R. Maire in the herbarium of the University Montpellier (MPU). Online available under https://science.mnhn.fr/institution/um/collection/mpu/item/mpu001465a.

**Electr. Suppl.: Fig. S5.**
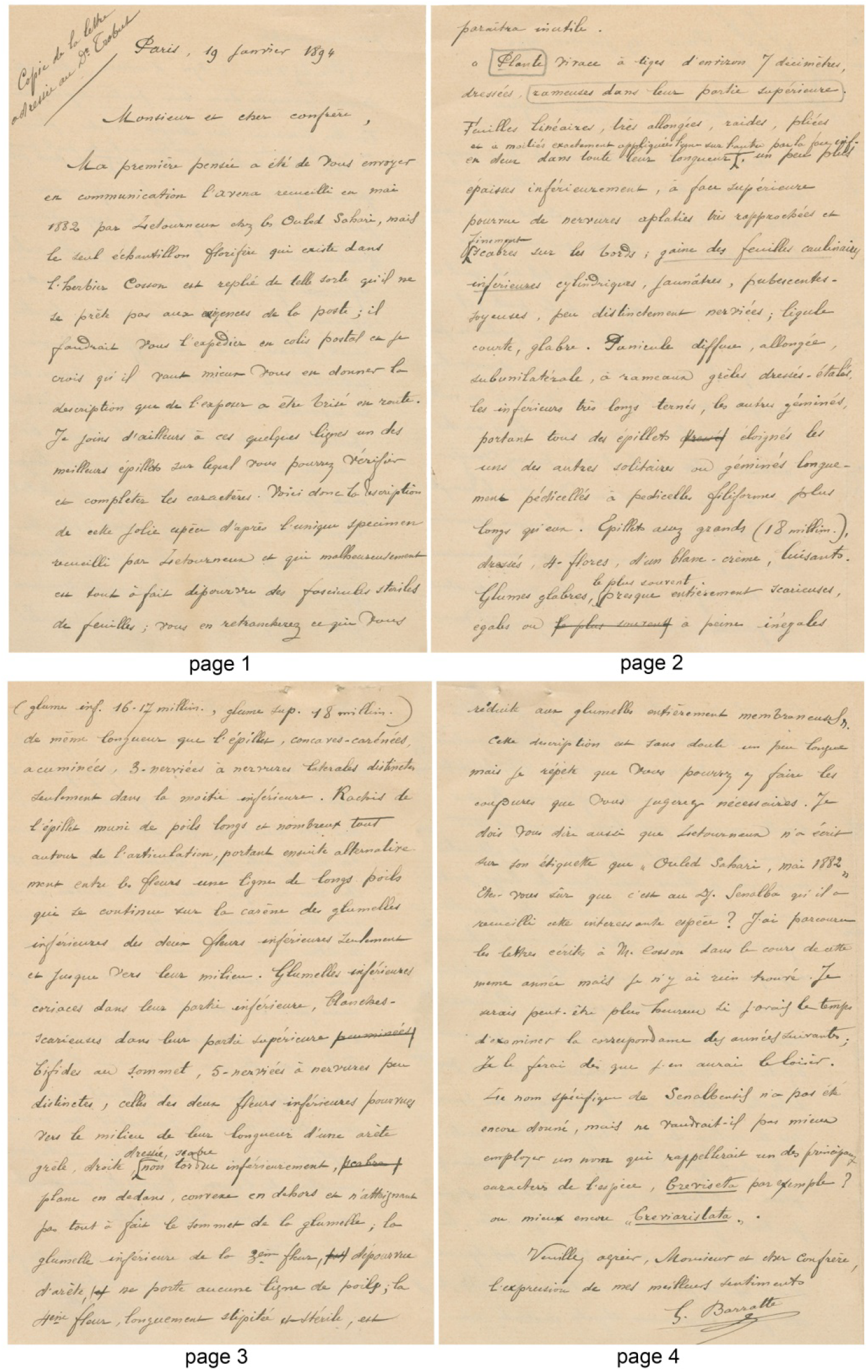
Barratte’s handwritten transcript of the letter he sent to Trabut. The letter is attached by a needle at the top of the holotype herbarium sheet of *Avena breviaristata* (P00152040). This letter was temporarily removed for Fig. 1 and for the online images of this voucher on the web pages of P and *JSTOR Global Plants* (see text). This “copie de la lettre adressée au Dr. Trabut” actually is the draft of the letter Barratte sent to Trabut, which is imaged in Fig. 3. See text for further explanation.

**Electr. Suppl.: Fig. S6.**
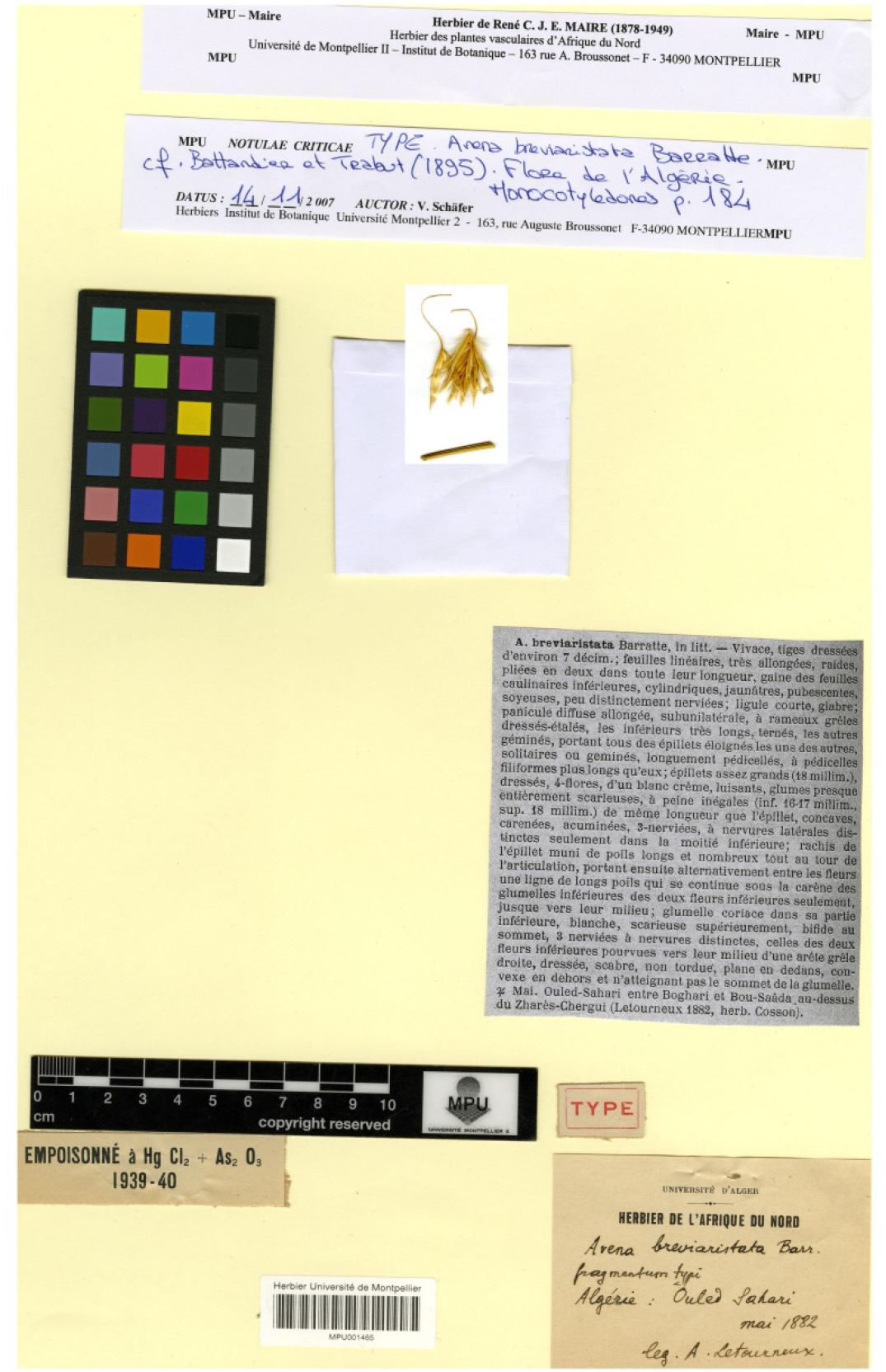
Fragment of the holotype (two spikelets) of *Avena breviaristata* in the herbarium of R. Maire deposited at MPU (voucher MPU001465).

**Electronic Supplementary Appendix S1.** Our transcript of Barratte’s original handwritten letter sent to Trabut preserved in the collection of R. Maire at the herbarium of the University Montpellier (MPU). Images of the letter are available online under https://science.mnhn.fr/institution/um/collection/mpu/item/mpu001465. Due to its considerable importance and the historical information on the type collection this letter is reproduced as Electr. Suppl. Fig. S5.

“Paris, 19 janvier 1894. Monsieur et cher confrère, Ma première pensée a été de vous envoyer en communication l’*Avena* recueilli en mai 1882 par Letourneux chez les Ouled Sahari, mais le seul échantillon florifère qui existe dans l’herbier est replié de telle sorte qu’il ne se prête pas aux exigences de la poste; il faudrait vous l’expédier comme colis postal et je crois qu’il vaut mieux vous en donner la description que de l’exposer a être brisé en route. Je joins d’ailleurs à ces quelques lignes un des meilleurs épillets sur lequel vous pourrez vérifier et compléter les caractères. Voici donc la description de cette jolie espèce d’après l’unique specimen recueilli par Letourneux etqui est malheureusement dépourvue des fascicules steriles de feuilles; vous en retrancherez ce qui vous paraîtra inutile. «Plante vivace à tiges d’environ 7 décimètres, dressée, rameuses dans leur partie supérieure. Feuilles linéaires, très allongées, raides, pliées en deux dans toute leur longueur et à moitiés exactement appliquées l’une sur l’autre par la face inférieure, un peu plus épaisses inférieurement, à face supérieure pourvue de nervures aplaties très rapprochées et finement scabres sur les bords; gaîne des feuilles caulinaires inférieures cylindriques, jaunâtres, pubescentes, soyeuses, peu distinctement nerviées; ligule courte, glabre. Panicule diffuse, allongée, subunilatérale, à rameaux grêles dressés-étalés, les inférieurs très longs ternés, les autres géminés, portant tous des épillets éloignés les uns des autres, solitaires ou géminés, longuement pedicellés à pédicelles filiformes plus longs qu’eux. Épillets assez grands (18 millim.), dressés, 4-flores, d’un blanc-crème, luisants. Glumes glabres, le plus souvent presque entièrement scarieuses, égales ou à peine inégales (glume inf. 16-17 millim., glume sup. 18 millim.), de même longueur que l’épillet, concaves-carénées, acuminées, 3-nerviées à nervures latérales distinctes seulement dans la moitié inférieure. Rachis de l’épillet muni des poils longs et nombreux tout autour de l’articulation, portant ensuite alternativement entre les fleurs une ligne de longs poils qui se continue sur la carène des glumelles inférieures des deux fleurs inférieures seulement et jusque vers leur milieu. Glumelles inférieures coriaces dans leur partie inférieure, blanches-scarieuses dans leur partie supérieure, bifides au sommet, 5-nerviées à nervures peu distinctes, celles des deux fleurs inférieures pourvues vers le milieu de leurs longueur d’une arête grêle, droite, dressée, scabre, non tordue inférieurement, plan en dedans, convexe en dehors et n’atteignant par tout à fait le sommet de la glumelle; la glumelle inférieure de la 3^ème^ fleur, dépourvue d’arête, ne porte aucune ligne de poils; la 4^ème^ fleur longuement stipitée et stérile est réduite aux glumelles entiérement membraneuses.» Cette description est sans doute un peu longue mais je répete que vous pourrez y faire les coupures que vous jugerez nécessaires. Je dois vous dire aussi que Letourneux n’a écrit sur son étiquette que «Ouled Sahari». Étes-vous sûre que c’est au Dj. Senalba qu’il [probably: où il] a recueilli cette intéressante espèce? J’ai parcouru les lettres écrites à M. Cosson dans le cours de cette même année mais je n’y ai rien trouvé. Je serais peut-être plus heureux si j’avais le temps de voir la correspondance des années suivantes; je le ferai dès que j’en aurai le loisir. Le nom spécifique de *Senalbensis* n’a certainement pas été donné a une espèce d’*Avena*, mais ne vaudrait-il pas mieux donner à cette jolie espèce un nom qui rappellerait un de ses principaux caractères, *breviseta* ou mieux encore *breviaristata*? Veuillez, Monsieur et cher confrère, agréer l’expression de mes meilleurs sentiments. G. Barratte”.

